# A simplified intermittent fasting regimen robustly extends *C. elegans* lifespan without FUdR or antibiotic confounds

**DOI:** 10.64898/2026.07.31.742121

**Authors:** Purbasha Dasgupta, Carlos G. Silva-García

## Abstract

Fasting-based dietary interventions are conserved regulators of aging that extend lifespan across species, including *Caenorhabditis elegans*. However, fasting studies in *C. elegans* are sensitive to experimental variables that can independently influence lifespan and health, including FUdR, antibiotic treatment, germline-less mutants, and the use of UV- or heat-killed bacteria. FUdR can alter lifespan, age-associated pathology, and stress responses, while antibiotics used to prevent bacterial growth during fasting may directly affect worm physiology. To minimize these confounding factors, we developed a simple adult-onset intermittent fasting paradigm that does not require FUdR, antibiotics, or bacterial killing. Wild-type worms were subjected to daily fasting periods of 5 h, 6 h, or 18 h until day 10 of adulthood and compared with continuously fed controls. Daily intermittent fasting robustly extended lifespan by 24-57%, demonstrating that repeated fasting windows during adulthood are sufficient to promote longevity under minimally confounded conditions. These findings establish a straightforward and experimentally tractable intermittent fasting paradigm for *C. elegans* and underscore the importance of limiting pharmacological and microbial conditions in dietary-intervention experiments.

## MAIN TEXT

Dietary interventions are potent regulators of aging that can modulate lifespan across multiple species, including humans (Partridge et al., 2005; Kapahi et al., 2017; Green et al., 2022; Schmauck-Medina et al., 2026). *Caenorhabditis elegans* has emerged as a powerful model for such studies, owing to its short lifespan, conserved longevity pathways, and uniform single-bacterial diet, which permits precise nutritional manipulation while minimizing the behavioral and social complexities present in mammalian systems (Lee et al., 2006; Honjoh et al., 2009; Pang & Curran, 2014). Among dietary interventions, fasting-refeeding approaches, including intermittent fasting, temporary fasting, and time-restricted feeding, are particularly robust drivers of longevity through evolutionarily conserved mechanisms (Uno et al., 2013; de Cabo & Mattson, 2019; Ivimey-Cook et al., 2021; Longo et al. 2021; Tatge et al., 2026). However, standard fasting protocols in *C. elegans* frequently rely on experimental manipulations that independently influence lifespan and complicate the interpretation of dietary effects.

Prolonged food deprivation promotes matricidal hatching in reproductively active hermaphrodites, often necessitating treatment with 5-fluoro-2′-deoxyuridine FUdR. Yet FUdR independently alters stress resistance, age-associated pathology, and lifespan in a context-dependent manner and can interact with environmental stressors to further extend lifespan (Anderson et al., 2016; Wang et al., 2019). Because fasting is inherently a stress-responsive intervention, combining it with FUdR may obscure the specific contributions of fasting-induced mechanisms. Additional experimental variables arise from efforts to suppress bacterial proliferation on fasting plates. To prevent worms from seeding food-free agar with gut-retained or cuticle-bound *E. coli*, studies commonly employ bactericidal antibiotics (e.g., streptomycin or carbenicillin), UV- or heat-killed bacteria, or peptone-depleted media **(Table 1)**. These approaches can independently affect longevity pathways. Antibiotics can extend lifespan by inducing mitonuclear protein imbalance and activating the mitochondrial unfolded protein response (UPR^mt^) (Houtkooper et al., 2013; Bonuccelli et al., 2023), whereas peptone depletion activates energy-sensing pathways, including AMPK and DAF-16/FoxO (Greer & Brunet, 2009; Stastna et al. 2015). Consequently, the lifespan extension observed under these conditions may reflect responses to pharmacologically or nutritional perturbations rather than physiological effects of fasting-refeeding cycles per se.

**Table 1:** Comparison of different fasting paradigms and their lifespan outcomes in C. elegans.

| Fasting regimen | Experimental conditions | Effects on longevity/mortality | References |
| --- | --- | --- | --- |
| Complete food removal (DR-FD) initiated day 2 of adulthood | FUdR (50 μM) on all plates; UV-killed OP50 for fed controls; N2 background | ~50% increase in median lifespan | Kaeberlein et al. (2006) |
| Complete dietary deprivation (DD); initiated at day 2 of adulthood for optimal effect (also tested at days 4 and 8); worms remain food-deprived for the remainder of their lives | FUdR (250 μg/mL), primarily fem-1(hc17) temperature-sensitive sterile mutant background | Up to 42.5% mean lifespan extension (day 2 DD initiation in sterile adults); detrimental when imposed on reproductively active day 0 adults | Lee et al. (2006) |
| Chronic bacterial deprivation (BD); complete removal of OP50 from NGM plates initiated at day 4 of adulthood; animals remain food-deprived for life. Also tested: BD initiated at days 8, 14, 20, and 24; and transient BD (day 2 to day 8, 14, or 28) with return to fed conditions | FUdR (50 μM); UV-killed OP50 on control-fed plates; N2 background | ~48-67% median lifespan extension (day 4 BD onset, N2); significant extension even when BD initiated after >50% of cohort had died (day 24); transient BD conferred lasting survival and thermotolerance benefits after return to fed diet | Smith et al. (2008) |
| Chronic bacterial deprivation from day 4 of adulthood | UV-killed OP50 on control plates; FUdR 50 μM | Mean LS +30%; max LS +30% (N2); wild isolates +23–51% | Sutphin & Kaeberlein (2008) |
| Alternate-day fasting (24h fast / 24h refed) or every-2-day fasting (48h fast / 24h refed) | FUdR (200 μg/mL), added at day 3 post-hatching, UV-killed OP50 was used for both AL controls and refeeding phases (not live bacteria); N2 background | 40.4% (alternate-day) and 56.6% (every-2-day) mean lifespan extension | Honjoh et al. (2009) |
| 2-day bacterial deprivation; late-L4 to day 2 of adulthood; returned to AL from day 3 | No FUdR, live OP50-1; no peptone on fasting plates; N2 background | log-odds = -1.69 (reduced mortality) | Ivimey-Cook et al. (2021) |
| Intermittent fasting: 4 h fasting/day for 5 consecutive days (days 1-5 of adulthood); refed on OP50 after each fast | FUdR (10 μM, post-fasting lifespan plates), Live OP50; N2 background | ~27% (4 h); ~25% (8 h); ~34% (12 h) | Li et al. (2026) |
| Single 24h fast at day 1 of adulthood; refed thereafter on standard NGM | FUdR (50 mM as reported), live OP50-1; N2 background | ~11% – 67% median lifespan extension across replicates | Zhou et al. (2026) |
| Single 24h fast at day 1 of adulthood; refed thereafter | FUdR (100 μg/mL); carbenicillin (100 μg/mL); IPTG (1 mM); HT115 bacteria | 40.8% median lifespan extension | Tatge et al. (2026) |
| 5 h and 6 h daily fasting initiated on day 1 of adulthood; 18 h overnight fasting initiated on day 3 of adulthood (post attainment of reproductive peak); all regimens maintained through day 10. | No FUdR, live OP50-1; no antibiotics or other bacteria-killing agents; N2 background | 57% (5 h IF), 24% (6 h IF), and 35% (18 h IF) | This study |

To minimize experimental confounders and build upon our previous studies of short-term fasting (Silva-Garcia et al., 2013; Lascarez-Lagunas et al., 2014; Huelgas-Morales et al., 2016), we developed an adult-onset intermittent fasting regimen that requires no FUdR, antibiotics, or other bacterial killing agents, on either fasting or fed plates. We tested three intermittent fasting protocols: 5 h and 6 h daily fasting initiated on day 1 of adulthood, and 18 h fasting initiated on day 3 of adulthood, after animals had reached their reproductive peak. All three regimens significantly extended median lifespan relative to *ad libitum* controls (5 h: 57% extension; 6 h: 24%; 18 h: 35%; log-rank p < 0.0001 for 5 h and 18 h, p = 0.0011 for 6 h; **Figure 1A-C**).

**Figure 1.**
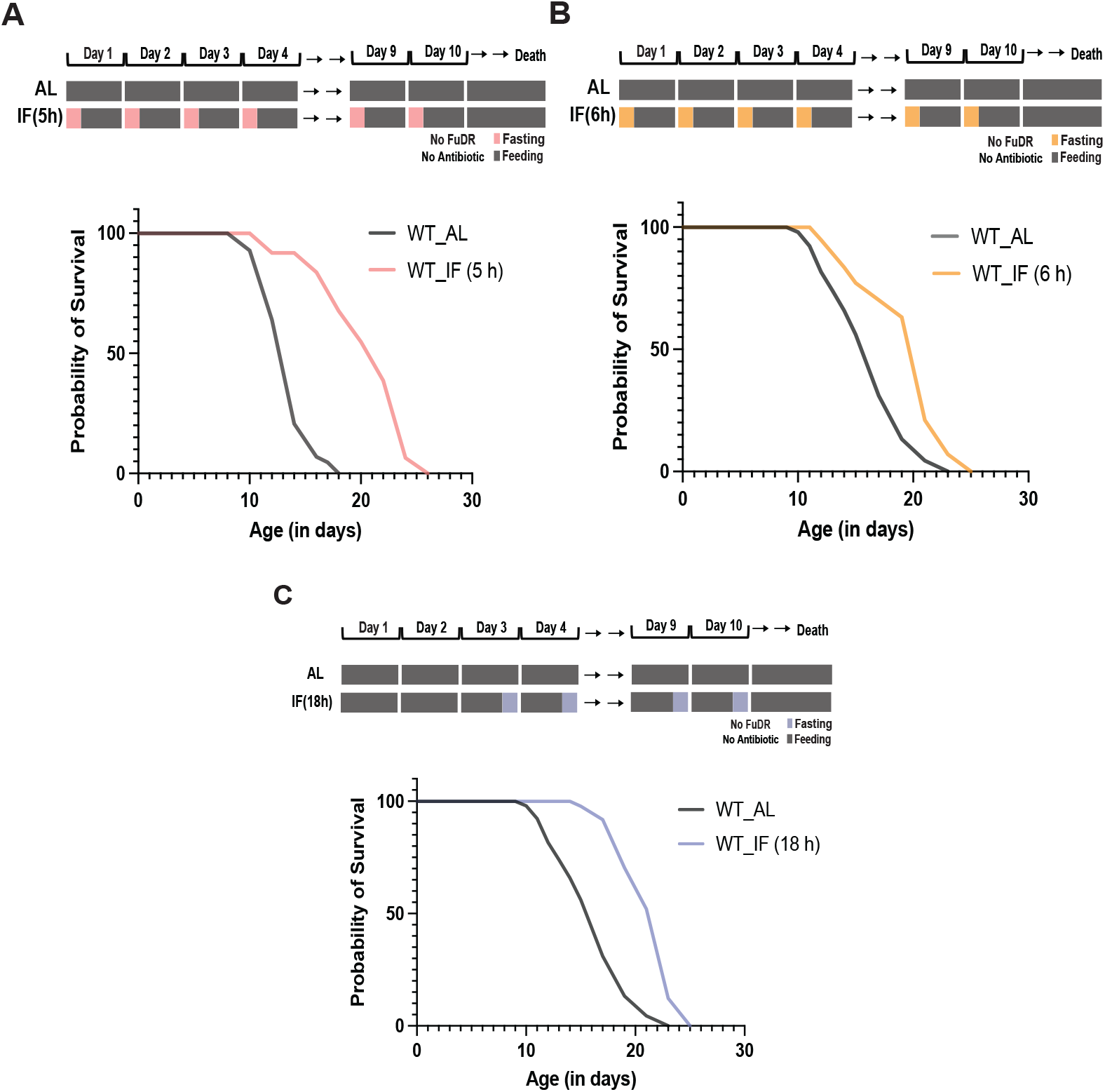
Adult-onset intermittent fasting extends C. elegans lifespan independently of FUdR and antibiotics. (A–C) Schematics (top) and survival curves (bottom) of N2 wild-type worms under intermittent fasting (IF) regimens compared to ad libitum (AL) controls. Daily 5 h (A) and 6 h (B) fasting was initiated on day 1 of adulthood; 18 h overnight fasting (C) was initiated on day 3 after the reproductive peak. AL controls were maintained on seeded plates throughout. All regimens continued through day 10. No FUdR, antibiotics, or bacterial killing agents were used. Median lifespan was extended by 57% (5 h IF; p < 0.0001), 24% (6 h IF; p = 0.0011), and 35% (18 h IF; p < 0.0001) relative to AL controls (log-rank test)

Together, these results demonstrate that short daily fasting bouts, as brief as 5 h, are sufficient to significantly extend lifespan under pharmacologically clean conditions. These findings are consistent with our previous work showing that short-term fasting profoundly remodels germline physiology by regulating apoptosis, stress granule formation, and protein translation (Silva-Garcia et al., 2013; Lascarez-Lagunas et al., 2014; Huelgas-Morales et al., 2016), highlighting the remarkable capacity of brief fasting periods to elicit broad physiological responses. Importantly, the longevity benefits observed in our study are comparable to those reported for established fasting-based interventions in *C. elegans* (**Table 1**).

Among the studies mentioned in Table 1, Li et al. (2026) is particularly noteworthy because its experimental design most closely resembles ours, employing short daily fasting windows in day 1 adults with live OP50 and reporting a modest but significant lifespan extension (25-34%). However, Li et al. acknowledge that the inclusion of FUdR limits their conclusions to relative lifespan comparisons performed under identical conditions. By contrast, our intermittent fasting paradigm is conducted entirely without FUdR, antibiotics, or other bacterial killing steps, allowing the longevity effects of fasting-refeeding cycles to be evaluated independently of pharmacological artifacts. Ongoing studies aim to define the molecular mechanisms underlying this response and to determine whether they overlap with or are distinct from those engaged by other fasting paradigms. Going forward, establishing the extent to which these mechanisms are conserved across species, including humans, will be critical for assessing their translational potential.

## METHODS

### *C. elegans* strains and husbandry

Wild-type N2 animals were obtained from the Caenorhabditis Genetics Center (CGC). The strain was maintained on standard nematode growth media (NGM) seeded with *E. coli* OP50-1 at 20°C.

### Bacterial strains

*E. coli* OP50-1 was cultured overnight in LB broth containing 50 μg/mL streptomycin at 37°C. For seeded plates, 100 μL of liquid culture was spread onto 60 mm NGM plates and allowed to grow for two days at room temperature prior to use.

### Intermittent fasting regimen

Synchronized populations were obtained by bleaching gravid adults. Animals were raised on 60 mm seeded NGM plates without FUdR until day 1 of adulthood. Three daily fasting durations were tested: 5 h, and 6 h initiated at day 1 of adulthood, and 18 h initiated at day 3 (post-reproductive peak). Prior to each transfer, animals were individually picked onto an adjacent unseeded agar surface or a separate unseeded NGM plate and allowed to crawl briefly to shed residual bacteria from the cuticle. Then, animals were transferred on 100 mm unseeded NGM fasting plates containing no FUdR, antibiotics, or other bacterial killing agents. The 100 mm plate format was used to reduce animal loss from edge-crawling; plates were inspected for agar cracks prior to use. After each fasting interval, animals were transferred to freshly seeded 60 mm NGM plates using a small amount of bacteria from the destination plates as a cushion to minimize mechanical stress during transfer. This cycle was maintained through day 10 of adulthood. *Ad libitum* (AL) controls were maintained continuously on seeded plates without FUdR and subjected to identical handling to account for transfer effects.

### Survival analyses

Lifespan assays were conducted at 20°C. Survival was scored daily through day 10, then every other day thereafter until death. Animals were considered dead upon failure to respond to three consecutive gentle prods with a platinum wire. Animals were censored for crawling off plates, bagging, vulval rupture, or plate contamination. To compensate for elevated censoring associated with intermittent fasting, cohort sizes were set at 300 worms for fasting conditions and 200 worms for *ad libitum* controls. Survival curves were compared using the log-rank (Mantel-Cox) method in GraphPad Prism.

## Acknowledgments

We thank all members of the Silva-García laboratory for their helpful discussions. Bristol N2 strain was provided by the Caenorhabditis Genetics Center (CGC), which is funded by the NIH Office of Research Infrastructure Programs (P40 OD010440).

